# COVID-19 and silent hypoxemia in a minimal closed-loop model of the respiratory rhythm generator

**DOI:** 10.1101/2023.04.19.536507

**Authors:** Casey O. Diekman, Peter J. Thomas, Christopher G. Wilson

## Abstract

Silent hypoxemia, or ‘happy hypoxia’, is a puzzling phenomenon in which patients who have contracted COVID-19 exhibit very low oxygen saturation (SaO_2_ < 80%) but do not experience discomfort in breathing. The mechanism by which this blunted response to hypoxia occurs is unknown. We have previously shown that a computational model (Diekman et al., 2017, J. Neurophysiol) of the respiratory neural network can be used to test hypotheses focused on changes in chemosensory inputs to the central pattern generator (CPG). We hypothesize that altered chemosensory function at the level of the carotid bodies and/or the *nucleus tractus solitarii* are responsible for the blunted response to hypoxia. Here, we use our model to explore this hypothesis by altering the properties of the gain function representing oxygen sensing inputs to the CPG. We then vary other parameters in the model and show that oxygen carrying capacity is the most salient factor for producing silent hypoxemia. We call for clinicians to measure hematocrit as a clinical index of altered physiology in response to COVID-19 infection.

## Introduction

### COVID-19 and silent hypoxemia

The global COVID-19 pandemic has led to over 1,000,000 deaths in the United States, and over 6,840,000 worldwide, since its onset in late 2019 [1]. COVID-19 can cause profoundly low levels of oxygen in the blood (hypoxemia) with near normal arterial carbon dioxide (*P*_a_CO_2_) levels. Although some individuals with COVID-19-induced hypoxemia experience dyspnea (breathing discomfort), many do not [2]. During surges of the pandemic, patients arriving at already overcrowded emergency rooms (ERs) who were not in obvious respiratory distress were often triaged [2]. However, some of these patients may have reduced oxygen saturation despite their lack of dyspnea [3, 4, 5]. This subpopulation of COVID-19 patients present with a novel condition known as *silent hypoxemia* or “happy hypoxia” [3]. Silent hypoxemia can result in tachypnea (rapid, shallow breathing), and with severe hypoxemia, changes in ventilation can occur [6, 7], but in general there is an absence of increased alveolar ventilation [2].

The mechanism underlying this condition is poorly understood but has been hypothesized to depend upon high expression levels of angiotensin converting enzyme 2 (ACE2) in the lungs, carotid body, and, perhaps, in the central breathing control circuitry within the medulla oblongata [3]. Additionally, recent work has shown that there is a shift in the oxyhemoglobin dissociation curve in COVID-19 patients [8, 9]. Since carotid body chemoreceptors respond to both low O_2_ and high CO_2_, a primary problem in these patients may be dysregulation of these sensors and chemosensory reflexes in general. ACE2 is the cellular entry point for SARS-CoV-2 [10]. COVID-19 infection has been shown to increase ACE2 expression, leading to changes in sensitivity to both CO_2_ and O_2_; changes in blood gases lead to a concomitant change in activity within the *nucleus tractus solitarii* (NTS). Recent work has shown that ACE2 is present within the carotid bodies of humans [11, 12] and there is evidence of altered chemosensation across multiple systems with SARS-CoV-2 infection [13]. The absence of dyspnea—even though patients exhibit low oxygen saturation—suggests that changes in carotid body inputs to the NTS are a key feature of SARS-CoV-2 infection. Additionally, there may be changes in NTS activity that contribute to the blunted ventilatory response but this has not yet been reported.

Given the low partial pressure of oxygen in arterial blood (*P*_a_O_2_) of patients infected with SARS-CoV-2 virus [14, 15] and the high expression of ACE2 in the carotid bodies, it is likely that altered chemosensory reflexes play a central role in the symptoms and outcomes seen in COVID-19 patients [11, 16]. In light of this data, we hypothesize that altered chemosensory function at the level of the carotid bodies and/or the NTS are responsible for this blunted response to hypoxia. We use a previously published computational model of respiratory control [17] to explore this hypothesis by altering the properties of the gain function representing oxygen sensing inputs to the respiratory central pattern generator (CPG). ^1^

### Quantitative modeling approach

Quantitative modeling has helped elucidate principles of normal and pathological functioning of the respiratory system, although its fundamental mechanisms remain debated. Mathematical models can be particularly helpful for generating experimentally testable hypotheses. A variety of models have been developed for the respiratory CPG [18, 19, 20, 21, 22, 23, 24, 25], for chemosensory feedback-based regulation schemes [26, 27, 28], and for cardiopulmonary gas exchange [29]. See [30, 31] for a review. A smaller number of published models represent closed-loop control incorporating a conductance-based CPG, muscle dynamics, gas exchange, and sensory feedback [32, 33, 34]. Of these, several focus on hypercapnia (excessive CO_2_) as the regulatory pathway. In order to generate hypotheses about silent hypoxemia, we chose to work with a conductance-based CPG model with O_2_ chemosensation as the sensory feedback pathway closing the control loop. To our knowledge, our previously published model [17] is the only model meeting these criteria. Aspects of it have been experimentally validated [35, 36]. Like any model, this model fails to represent all aspects of the control system. We have not included CO_2_ sensing in our model due to the high diffusion rates of CO_2_ when compared to O_2_ in the lung [37] and evidence showing that CO_2_ is *≤* 35 mmHg in patients presenting with silent hypoxemia (SH) and minimal tachypnea [5, 38] Additionally, we do not explicitly include rapidly adapting (RAR) or slowly adapting (SAR) lung mechanoreceptors in the model—lung volume is present in the model and reproduces inspiratory drive in much the same way that SARs do in vivo. Nevertheless, in spite of these limitations, the model suffices to generate testable hypotheses that could be pursued by the clinical community.

The model studied in [17] has seven dynamical variables: voltage of a central pacemaker cell, together with one fast and one slow gating variable; diaphragm muscle activation; lung volume; partial pressure of O_2_ in the lung; and partial pressure of O_2_ in the bloodstream. Regulation of the endogenous breathing rhythm occurs through hypoxia-sensitive chemosensory feedback in the model. Thus we will refer to this system as the 7D-O2 model. Figure 1 shows the closed-loop structure of the 7D-O2 model. We briefly describe the 7D-O2 model below. For a full description of the nonlinear system of seven ordinary differential equations specifying the model see the Methods section.

**Figure 1:**
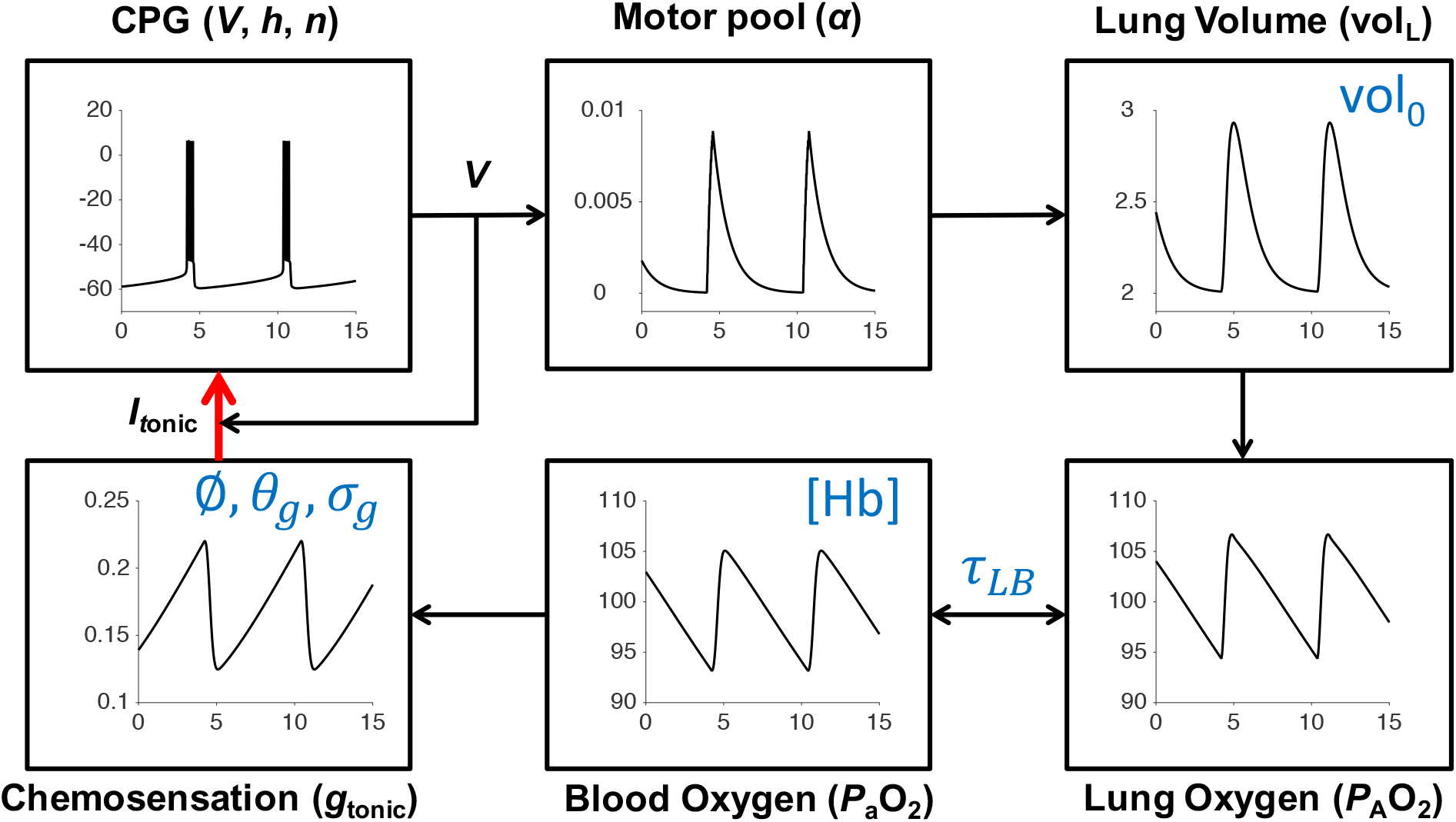
Schematic of the 7D-O2 model. Bursts of action potential firing (*V*, mV) in the respiratory central pattern generator (CPG) drive a pool of motor neurons (*α*, dimensionless) leading to expansions of lung volume (vol_L_, L) and increases in lung and blood oxygen (*P*_A_O_2_ and *P*_a_O_2_, mmHg). Through a chemosensory pathway (*g*_tonic_, nS), the blood oxygen level affects the amount of excitatory current sent to the CPG, thereby closing the control loop (red arrow). Time (*t*, seconds) is the horizontal axis for all traces. The six parameters shown in blue are varied in this study to model silent hypoxemia. Redrawn, with modifications, from [17].

The 7D-O2 model is a closed-loop respiratory control model that comprises a well-established conductance-based central rhythm generator (the Butera-Rinzel-Smith model [18, 17]) with a voltage variable *V*, a fast gating variable (delayed-rectifier potassium current activation, *n*) and a slow gating variable (persistent sodium current inactivation, *h*). The output of the BRS model cell, namely the voltage, drives a motor pool activation variable, *α*, that in turn drives expansion of the lungs. The lung volume (vol_L_), the partial pressure of oxygen in the lungs (alveolar pressure, *P*_A_O_2_), and the partial pressure of oxygen in the bloodstream (*P*_a_O_2_) complete the model variables. The BRS cell includes an excitatory current driven by a tonic conductance that is regulated by chemosensory feedback, closing the control loop. The model includes a metabolic demand parameter, *M*, regulating the rate at which oxygen is removed from the bloodstream to the tissues. As the “phenotype” or “physiology” of the model, we take the steady-state value of *P*_a_O_2_ as a function of *M*. For the original model as presented in [17], the *P*_a_O_2_-vs-*M* curve shows a plateau near 100 mm Hg (normoxia) that collapses to a critically hypoxic state when *M* increases past a high threshold (Fig. 2A). As we varied the original parameters to investigate possible mechanisms of silent hypoxemia, we monitored the height of the normoxia plateau, and the location of the collapse point.

**Figure 2:**
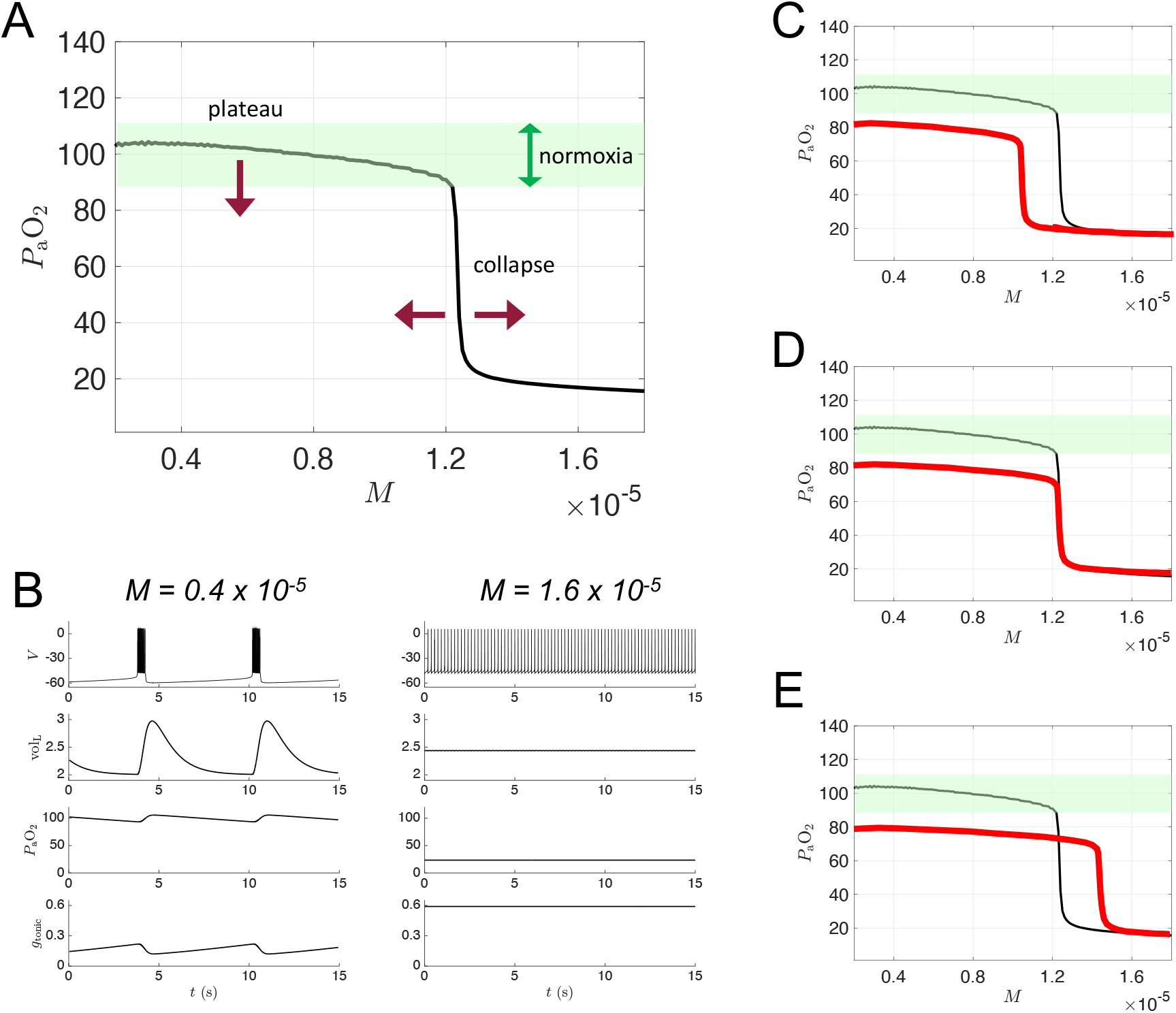
Dynamics of 7D-O2 model and hypothetical silent hypoxemia *P*_a_O_2_ vs *M* curves. **A:** Blood oxygen *P*_a_O_2_ as a function of metabolic demand (*M*) in the original 7D-O2 model. There is a plateau region at low *M* values for which normoxia (green shading) is maintained, and a collapse point at approximately *M* = 1.2 × 10^−5^ ms^−1^ beyond which severe hypoxia occurs. In a model of silent hypoxemia, it seems clear that the plateau portion of the curve should shift lower (maroon arrow pointing down), but it is not as clear whether the collapse point should remain in the same location or shift horizontally (maroon arrows pointing left and right). **B:** Variables of the 7D-O2 model for *M* values in the plateau region (left column) or after the collapse point (right column). **C-E:** Hypothetical silent hypoxemia models (red) with a lower plateau and a collapse point shifted to a lower *M* value (**C**), in the same location (**D**), or shifted to a higher *M* value (**E**) compared to the 7D-O2 model (black).

### Relating model parameters to potential silent hypoxemia mechanisms

The mechanism by which COVID-19 leads to sustained hypoxemia in the absence of dyspnea is currently unknown. The minimalist model of [17] includes a number of key parameters that are plausible targets for modification to mimic the effects of COVID-19-infection on respiratory control. After three years of the COVID-19 pandemic and on-going endemic infection, a few key patho-physiologies have become apparent. First, ACE2 expression is correlated with the location and severity of infection [39]. Because ACE2 is, based on current knowledge, the main vector by which SARS-CoV-2 enters the body’s cells, changes in ACE2 expression should have an impact on the severity and time course of COVID-19 symptoms. Second, changes in NTS signaling may play a key role in altering the normal, physiological response to COVID-19, and that information may be carried by the glossopharyngeal nerve (innervating the carotid body) or lung afferents via the vagus nerve. Information sensed at the carotid bodies (and lung interoceptors) ultimately reaches the *nucleus tractus solitarii* via the vagus and glossopharyngeal nerves. From the NTS, these signals are distributed to local visceral integration circuits within the medulla, including the cardivascular control regions (rostral and caudal in the ventral medulla) and the preBötzinger complex and associated regions of respiratory control within the brainstem. Based on the clinical observations reported so far, it appears that there is a change in gain in the pathway from carotid body, to NTS, to the breathing rhythm generator and pattern formation network. These observations in patients have provided the motivation for us to focus on assessing the effect of changes in sensitivity/gain in this signaling pathway. This change in gain may be more prevalent in any one of these circuit elements and further work needs to be done to determine the exact mechanism by which sensitivity of the control circuit is impacted.

Oxygen carrying capacity is a key variable in pulmonary mechanics. Repeated bouts of intermittent hypoxia, as seen in obstructive sleep apnea, can increase HIF-1*α* signaling, with a subsequent increase in erythropoietin, and an increase in hemoglobin and erythrocytes. Similar changes are seen in conditions that result in chronic hypoxemia and hypercapnia, such as cardiovascular disease, obstructive sleep apnea, and chronic obstructive pulmonary disease [40, 41, 42]. Many of the patients presenting with silent hypoxemia have pre-existing conditions and co-morbidities that are likely to increase hematocrit and this increase in oxygen carrying capacity may blunt chemoreceptor responses—exacerbating the “happy hypoxia” phenomenon. Unfortunately, no current literature quantifies hematocrit in these patients.

Motivated by these observations, we systematically varied (plus or minus 20%) the following parameters that control the saturating effect of hypoxia-sensitive chemosensory feedback to the central pattern generator: *σ*_g_, which controls the slope of the sensory feedback curve at maximum sensitivity (gain at threshold); *θ*_g_, which controls the threshold activation value for sensory feedback (50% activation point); and *φ*, which controls the maximum sensory feedback drive at full activation. Lung volume is a key determinant of mechanosensory feedback to the NTS and the CPG. Our model incorporates lung volume and allows us to monitor changes in lung volume in response to changes in central drive for breathing. This also allows us to monitor lung volume as an outcome measure to determine if the CPG is actually causing lung inflation in a way that assures sufficient gas exchange to sustain life when extrapolated to animal models or human subjects.

Ventilation-perfusion matching is a key drive for respiration. In mammals the interplay between cardiovascular and respiratory control is essential for ensuring that sufficient oxygen is delivered to the body and CO_2_ is removed via the lung. We have included a time constant for O_2_ transport between the lung and blood which allows us to simulate changes in diffusion and dwell time within the lung that correlate with diseases such as chronic obstructive pulmonary disease (COPD) and lung fibrosis. Oxygen consumption and CO_2_ production are key elements for determining how changes in breathing can match metabolic demand. We have included a simplified treatment of metabolism in the model. As a “biomarker” to test the model behavior, for all parameter sets we varied the metabolic demand parameter *M* across a range of values. We have not included CO_2_ in this model, because CO_2_ diffuses up to 20 times faster than O_2_ [37] and patients with SH do not appear to be hypercapnic since there is very little change in breathing rate—CO_2_ is a potent stimulator of minute ventilation and hypercapnia results in pronounced increases in breathing frequency [43, 44, 45].

Additionally, we vary the hemoglobin concentration to mimic the effect of chronic hypoxia seen in humans living in hypoxic environments which can include mountain dwellers [46], individuals with severe obstructive sleep apnea [42], or other cardio-respiratory disorders [41, 47]. These individuals can have high hematocrit, a corresponding increase in red blood cells, and increased blood viscosity—similar to what has been reported in COVID-19 patients [48].

## Results

Figure 1 shows a schematic of the respiratory control model that we will use in this study, with components representing CPG membrane potential (*V*), motor pool activity (*α*), lung volume (vol_L_), lung oxygen (*P*_A_O_2_), blood oxygen (*P*_a_O_2_), and chemosensation (*g*_tonic_). The model has a closed-loop structure since an excitatory current, *I*_*tonic*_, depends on *P*_a_O_2_ and is an input to the CPG component (red arrow). In this model, the rate of metabolic demand for oxygen from the tissues is represented by the parameter *M*. If metabolic demand is low or moderate (*M* < 1.2 × 10^−5^ ms^−1^), then the model exhibits a stable eupneic rhythm with CPG bursting activity driving fluctuations in lung volume that bring in a sufficient amount of oxygen to maintain *P*_a_O_2_ in the normoxia range (see the “plateau” region of the *P*_a_O_2_ versus *M* curve shown in Fig. 2A and the traces in the left panel of Fig. 2B.) However, if metabolic demand is too high (*M >* 1.2 × 10^−5^), then the model exhibits a form of tachypnea, where CPG bursting activity is replaced with tonic spiking that does not drive the lungs effectively enough to maintain *P*_a_O_2_ in the normoxia range (see the “collapse” region of Fig. 2A and the right panel of Fig. 2B).

In silent hypoxemia, we would expect to observe a lower height for the plateau region of the *P*_a_O_2_ versus *M* curve, since these patients display abnormally low *P*_a_O_2_ despite minimal changes in minute ventilation. There are three possibilities regarding the collapse region in silent hypoxemia patients: the collapse point could shift to a lower *M* value (as illustrated in Fig. 2C), stay at the same *M* value (as in Fig. 2D), or shift to a higher *M* value (as in Fig. 2E). Because it seems plausible that a disease-induced reduction in steady-state *P*_a_O_2_ would be accompanied by an *decrease* in tolerance of higher metabolic demand, we explored parameter space to see if the closed-loop model is capable of producing *P*_a_O_2_ versus *M* curves with shapes similar to the hypothetical curve shown in Fig. 2C.

Motivated by the hypothesis that silent hypoxemia results from a dysregulation of carotid body O_2_ receptors, we first considered variation of the parameters associated with the chemosensory pathway of the model. In the 7D-O2 model, there is a sigmoidal relationship between *P*_a_O_2_ and *g*_tonic_, with the parameters *φ, θ*_*g*_, and *σ*_*g*_ controlling the height, half-activation, and slope of the sigmoid, respectively (see Fig. 3A). We simulated the closed-loop model over a range of *M* values while varying these parameters over 3 levels spanning roughly ±20% of their original values (*φ* = 0.24, 0.3, 0.36, *θ*_*g*_ = 70, 85, 100, and *σ*_*g*_ = 0.24, 0.3, 0.36), yielding 27 different combinations in total. Figure 3B shows that varying these parameters generates *P*_a_O_2_ vs *M* curves in which the plateau and collapse point are shifted down and to the right (similar to the hypothetical case shown in Fig. 2E). None of the 27 combinations, however, produce any curves with the plateau and the collapse point shifted down and to the left (similar to Fig. 2C). Thus, we do not consider any of these model variants to suitably capture the phenomenon of silent hypoxemia.

**Figure 3:**
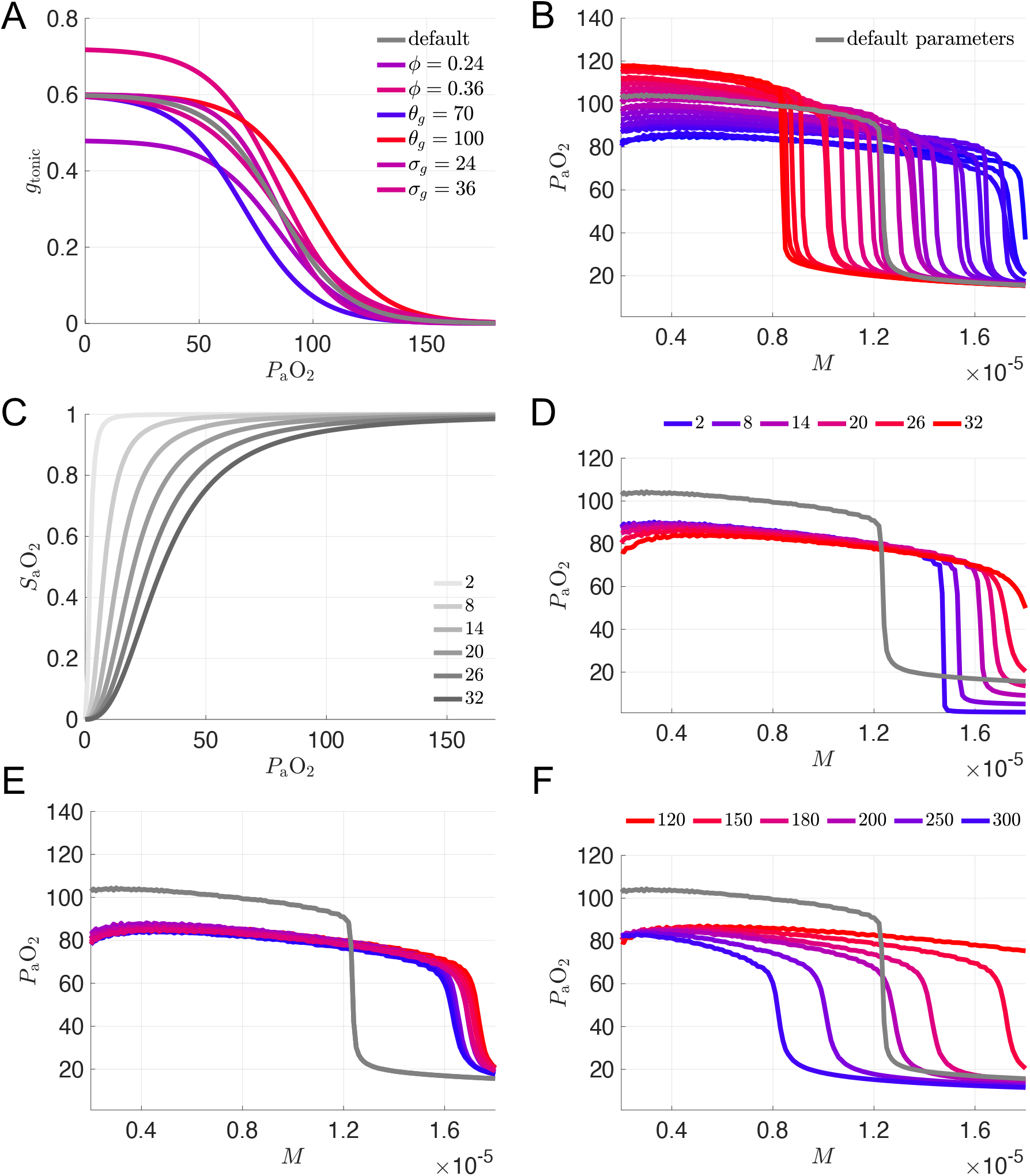
Sensitivity of *P*_a_O_2_ vs *M* curves to variation of model parameters. **A:** Chemosensory sigmoid of *g*_tonic_ as a function of *P*_a_O_2_ with various parameter values for the maximum (*φ*), half-activation (*θ*_*g*_), and slope (*σ*_*g*_) of the sigmoid. Default settings from the original 7D-O2 model (*φ* = 0.3 nS, *θ*_*g*_ = 85 mmHg, *σ*_*g*_ = 30 mmHg) shown in gray. See panel (B) for the definition of the color scale used for the other curves. **B:** *P*_a_O_2_ vs *M* curves for 27 different combinations of the chemosensory sigmoid parameters (*φ* = 0.24, 0.3, 0.36; *θ*_*g*_ = 70, 85, 100; *σ*_*g*_ = 24, 30, 36) on a color scale with the lowest and highest maximum *P*_a_O_2_ values shown in blue and red, respectively, with the exception of the default parameter set which is shown in gray. **C:** Hemoglobin saturation curves (Eq. 18) for various hemoglobin binding affinities *K*. Default model has *K* = 26 mmHg. **D:** *P*_a_O_2_ vs *M* curves for the set of *K* values shown in (**C**). **E:** *P*_a_O_2_ vs *M* curves for 9 different combinations of oxygen flux and lung volume parameters (*τ*_LB_ = 100, 500, 900 ms; vol_0_ = 1.6, 2.0, 2.4 L), with a constant set of chemosensory sigmoid parameters (*φ* = 0.24, *θ*_*g*_ = 70, *σ*_*g*_ = 36) on a color scale with the lowest and highest *M* values at the collapse point (*P*_a_O_2_ = 40) shown in blue and red, respectively, with the exception of the default parameter set which is shown in gray. **F:** *P*_a_O_2_ vs *M* curves for 6 different values of hemoglobin concentration [Hb], with *τ*_LB_ = 500, vol_0_ = 2.0, and the same chemosensory sigmoid parameters and color scale as in (**E**). The purple [Hb] = 250 curve was selected as a putative model for silent hypoxemia.

**Figure 4:**
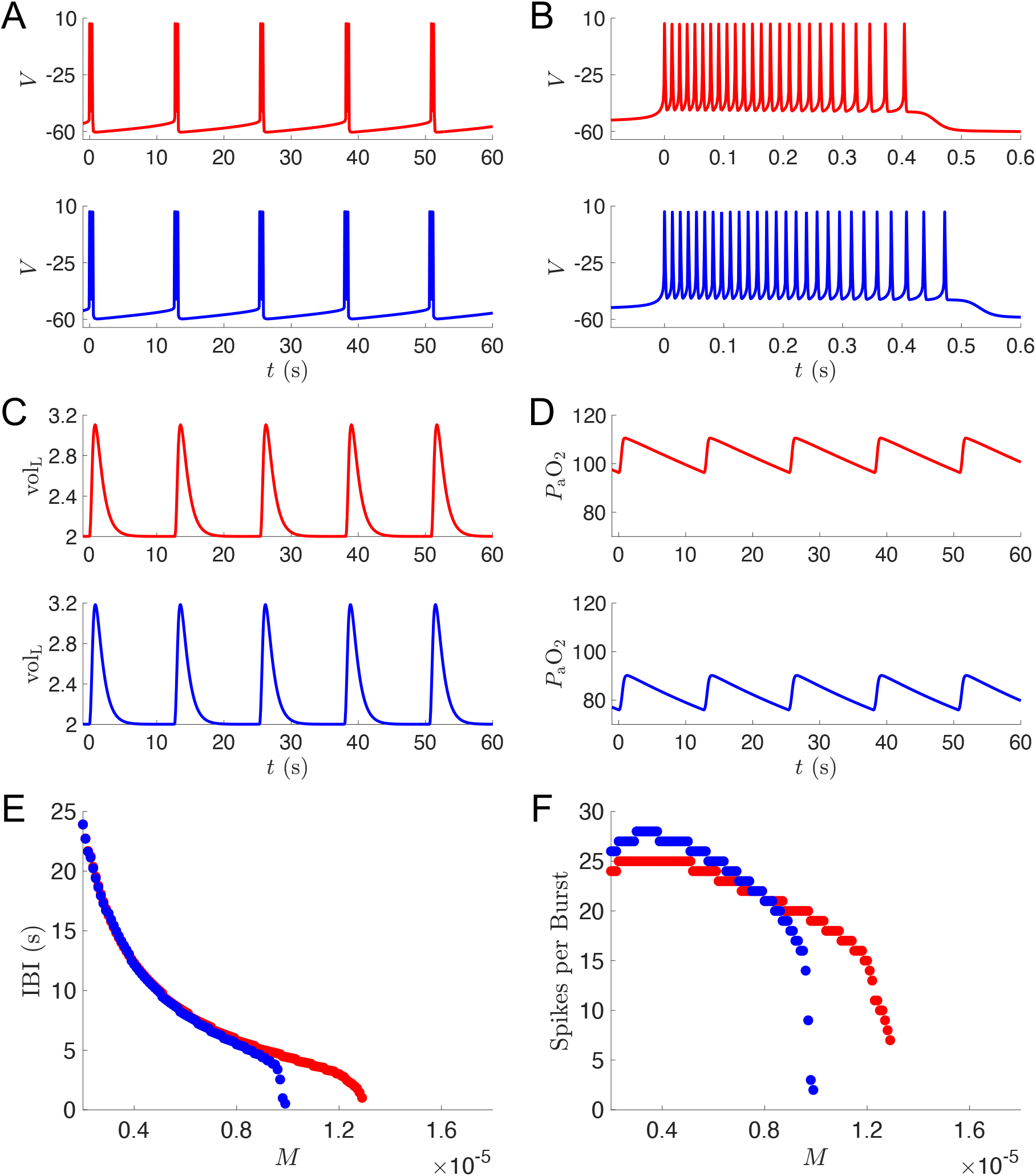
Simulations of putative silent hypoxemia model. **A-F:** Output from simulations of the normoxia model (the original 7D-O2 model) and the silent hypoxemia model ([Hb] = 250 curve from Fig. 3D) shown in red and blue, respectively. Voltage traces showing multiple bursts (**A**) and zooming in on a single burst (**B**); lung volume (**C**) and blood oxygen (**D**) traces across multiple bursts. For panels (**A-D**), *M* = 0.4 × 10^−5^ ms^−1^. Interburst interval (**E**) and number of spikes per burst (**F**) as a function of metabolic demand.

In order to proceed further, we selected a single parameter set from among the 27 combinations as our working model for producing the hypoxic plateau region, namely *φ* = 0.24, *θ*_*g*_ = 70, and *σ*_*g*_ = 36. These parameters gave the curve with the greatest reduction of *P*_a_O_2_ (darkest blue curves in Figs. 3A and B), although the collapse point did shift to significantly higher values of *M*.

Since varying the chemosensory parameters alone was not sufficient to model a silent hypoxemia patient prone to respiratory collapse, we considered other parameters that could plausibly be affected by COVID-19. Based on reports indicating that COVID-19 patients have altered oxyhemoglobin dissociation curves [8, 9], we considered variation of the model parameter *K* which represents hemoglobin binding affinity (Eq. 18 in Methods). The effect that increasing the binding affinity (decreasing *K*) has on the SaO_2_ *− P*_a_O_2_ saturation curve with the new chemosensory parameters is shown in Fig. 3C (see also Appendix Fig. 5A for the effect of varying *K* with the original chemosensory parameters). Tighter binding affinities (*K* values less than the default value of 26 mmHg) do shift the *P*_a_O_2_ vs *M* curve to the left, but the respiratory collapse point is still at higher metabolic demand values than the original model (Fig. 3D). Thus, we contined to explore other parameters that might plausibly be affected by COVID-19. For example, lung damage due to excessive immune response or local thrombosis could reduce the effective unloaded lung volume (model parameter vol_0_), or impede the flux of oxygen between the alveoli and the alveolar capillaries. The latter effect could be reflected by an increase in the model parameter *τ*_LB_, which governs the effective relaxation time for differences in partial pressure of oxygen in the model’s lung and blood compartments, respectively. Therefore, while keeping the chemosensory sigmoid parameters (*φ* = 0.24, *θ*_*g*_ = 70, *σ*_*g*_ = 36) fixed, we varied the unloaded lung volume (vol_0_ = 1.6, 2.0, 2.4) and the time constant for the flux of oxygen from the lung to the blood (*τ*_LB_ = 100, 500, 900).

As shown in Fig. 3E, varying vol_0_ by ±20% and varying *τ*_LB_ by ±400% had surprisingly little effect on the height of the *P*_a_O_2_ versus *M* plateau, and did not significantly affect the collapse point either.

Finally, we considered variation of the parameter [Hb] representing the hematocrit, i.e. the concentration of hemoglobin within the blood, which was set to 150 g/l in the original 7D-O2 model. Fig. 3F shows that *increasing* [Hb] within the model *lowers* the collapse threshold of the *P*_a_O_2_ versus *M* curve, while maintaning a hypoxemic plateau around 80 mmHg. A 33% increase in [Hb] shifts the collapse point to a similar *M* value as the original 7D-O2 model, consistent with the hypothetical silent hypoxemia *P*_a_O_2_ vs *M* curve shown in Fig. 2D. Further increases in [Hb] yield collapse points with even lower *M* values, consistent with the hypothetical silent hypoxemia *P*_a_O_2_ vs *M* curve shown in Fig. 2C. See also Appendix Fig. 5B for the effect of varying [Hb] with all other parameters set to their original 7D-O2 model values.

We next considered the model with [Hb]=250 (the second curve from the left in Fig. 3D) as a putative example of silent hypoxemia, and analyzed the model dynamics for simulations in the plateau region and in response to increases in metabolic demand. Figure 4A shows voltage traces in the plateau region (*M* = 0.4 × 10^−5^ ms^−1^) for both the silent hypoxemia model (blue) and the original 7D-O2 normoxia model (red). The frequency of bursting is similar in the two models, but there are a few more spikes per burst in the hypoxemia model (Fig. 4B). This leads to slightly more vigorous lung expansions in the hypoxemia model (Fig. 4C), however the levels of oxygen in the blood remain substantially lower (Fig. 4D). As the metabolic demand is increased, the frequency of bursting in the hypoxemia model becomes much faster than in the normoxia model (Fig. 4E), and there are substantially fewer spikes per burst (Fig. 4F). This type of bursting activity leads to more frequent but less vigorous lung expansions and ultimately respiratory collapse at lower levels of metabolic demand in the hypoxemia model compared to the normoxia model (see Appendix Fig. 6).

## Discussion

We originally hypothesized that altered chemosensory input to the carotid bodies and, eventually, to the NTS and the rest of the breathing control circuitry, is a key factor in silent hypoxemia. However, our simulation results suggest that while changes in chemosensivity may play a role in silent hypoxemia, changes in metabolism and oxygen carrying capacity may have greater relevance for replicating the respiratory collapse seen in these patients. Specifically, altered chemosensitivity can create a hypoxemic plateau region (SaO_2_ < 90 mmHg) for a broad range of metabolic demand levels (*M* = 0.4 to 1.5×10^−5^ ms^−1^, see blue curves in Fig. 3B). When hemoglobin concentration is then increased, moderate levels of metabolic demand (*M* = 0.8 to 1.0×10^−5^ ms^−1^) lead to complete respiratory collapse (SaO_2_ < 60 mmHg, see blue and purple curves in Fig. 3F).

Our hypothesis was based on the premise that O_2_ sensing is the key factor in SH. Canonically, it has been suggested that CO_2_ is a primary driver for dyspnea [49, 50, 51], but there is evidence that both hypoxia and hypercapnia equivalently drive the sensation of air hunger [43]. However, clinical case and cohort studies show that patients with SH are not hypercapnic [5, 38]. This suggested to us that dysregulation of O_2_ sensation is a key contributor to the issues seen in SH. We tested this hypothesis by changing O_2_ sensitivity in the model at the level of the carotid bodies/NTS and evaulating whether those changes could reproduce the SH phenotype.

A complicating factor for these patients includes co-morbidities that have demonstrated correlation with poor outcome in patients with COVID-19. These comorbidities include obstructive sleep apnea (OSA), chronic obstructive pulmonary disease (COPD), or cardiovascular disease (including hypertension or heart failure). Patients suffering from these diseases often develop polycythemia— an increase in the hemoglobin and hematocrit to adaptively increase the O_2_ carrying capacity of the blood. High-altitude populations are well-adapted to chronic hypoxia and typically have a higher hematocrit in Andean populations versus Himalayan high-altitude dwellers [52], likely due to different adapation mechanisms. However, subjects with cardiovascular disease [53], obstructive apnea [54], and familial hyperlipidemia [41] also show increased hematocrit.

One consequence of pumping thicker blood is to increase the metabolic demand, even during rest. As we show in our results, increasing metabolic demand increases the likelihood of respiratory collapse. Somewhat paradoxically, with an increased oxygen carrying capacity, the patient may be less able to compensate for the worsening *P*_a_O_2_ and a critical tipping point for metabolic demand is reached where respiratory efforts are insufficient to keep up with demand. We have not yet seen any report documenting changes in hematocrit in COVID-19 patients who exhibit silent hypoxemia. Based on our modeling results, we would predict that these patients may show increased hematocrit levels. In support of our prediction, a recently published study [48] showed that higher blood viscosity was associated with an increase in mortality in COVID-19 patients. Obtaining this kind of data should be possible for patients admitted to the intensive care unit and should be a priority for future investigation.

Angiotensin-Converting Enzyme 2 (ACE2) is expressed in the lungs, carotid bodies, and respiratory region of the brainstem, and is likely the vector by which the SARS-CoV-2 virus invades the carotid bodies and/or the NTS, thereby potentially contributing to silent hypoxemia. High ACE2 levels also occur in the most vulnerable target organ systems seen in COVID-19 (elevated expression levels occur in lung, heart, ileum, kidney and bladder [39]). Since ACE2 expression is very high in the lungs, and since diffuse alveolar damage, bronchopneumonia, and alveolar hemorrhage are common in COVID-19 [40], it seems reasonable to hypothesize that the decrease in gas exchange across the alveolar membranes within the lung can alter not just the O_2_ carrying capacity but also increase metabolic demand for perfusion of the damaged lung. It may be of value to assess differences in mitochondrial activity in lung cells from normal and COVID-19 patients, or in animal models that have used SARS-CoV-2 or spike protein (now commercially available) to mimic the lung damage seen in human patients. Such experiments would provide data concerning cellular metabolism and give us greater understanding of the impact COVID-19 has on metabolic demand at all tissue levels.

Lack of dyspnea (breathing discomfort) in patients arriving at already overcrowded emergency rooms leads to triaging patients not in obvious respiratory distress, when in fact these patients often have reduced oxygen saturation [55]. Perhaps the greatest mystery that remains unresolved is why dyspnea is not typically seen in patients exhibiting silent hypoxemia. Sensory perception is subjective and can vary with a host of factors that include sex, socioeconomic background, and ethnicity [56, 57, 58]. There is some controversy about these correlates but they may be underlying factors that influence the reporting of silent hypoxemia. Once again, some demographic data is available concerning COVID-19 infection, mortality, and morbidity, but this information has not been correlated with silent hypoxemia yet. Ideally, demographic factors should be reported along with other patient data to better understand the incidence and severity of silent hypoxemia and dyspnea.

Patients with COVID-19 are also subject to mitochondrial dysregulation that contributes to severity and lethality. Mitochondrial function is impacted by the “cytokine storm”, a hallmark of the immune response to COVID-19. Thus, upregulation of cytokine release in the context of comorbidities that increase inflammation, including metabolic syndrome, obesity, type 2 diabetes, and increasing age—in addition to the lung and cardiovascular diseases mentioned above, are all associated with mitochondrial dysfunction [59, 60, 61]. SARS-CoV-2 infection causes multi-system changes at transcriptomic, proteomic, and metabolomic levels, altering normal cellular metabolism and changing mitochondrial respiration [61]. Disruption of normal mitochondrial function can result in an increase in reactive oxygen species (ROS) further exacerbating inflammation and increasing the likelihood of poor outcomes [62]. The “long COVID” phenomenon may be related to redox imbalance, which may be exacerbated by COVID-induced changes in mitochondria [63, 64] and, ultimately, fatigue related to metabolic impairment. Our results suggest that there is a delicate balance between metabolic demand changes and respiratory failure. One can easily speculate that reduction in available oxygen in concert with an increase in metabolic demand as the virus takes over cellular machinery to produce more viral particles can result in a point of critical failure.

One way to test this mechanism woud be to assay mitochondria function obtained in biopsies of tissue from COVID-19 patients or through animal models. Testing mitochondrial metabolism would be easier than using stress tests or cycle ergometry to determine metabolic load and ventilation-perfusion changes. Whole body tests would be problematic in COVID-19 patients and put them at greater risk for respiratory collapse. As long COVID has become better described, central nervous system (CNS) involvement and increased chronic inflammation are seen as sequelae that may continue to alter metabolism and mitochondrial function [65]. Further research is needed to determine if these effects are exacerbated by persistent metabolic impairment and whether symptoms such as cognitive fog depends on mitochondria and ROS handling problems within the CNS.

The relationship between changes in overall metabolic demand and cellular level metabolism has not yet been fully explored in COVID-19 patients. This is an important area for investigation, because while the metabolic demand required to pump more viscous blood [48] may selectively impact the cardiovascular system the most, metabolic demand may be increased systemically based on the diffuse organ involvement seen in these patients.

In addition to the limitations to our model that we have mentioned previously, we realize that our model represents a very reduced number of the elements in the central pattern generator and pattern formation network for breathing control. The brainstem network includes hundreds of neurons that participate in each breath [66, 67], and we have simplified this relatively complex circuit for the sake of rapid simulation time to test our hypotheses about SH. This heavily reductionist treatment of the brainstem network is an obvious limitation to simulation of the interacting populations of respiratory neurons and makes it difficult to interrogate the precise mechanisms by which respiratory collapse occurs in SH. Previously, we have demonstrated that increasing extracellular [K^+^] resulted in a progressive increase in respiratory rhythm that showed periodic, multi-periodic, quasi-periodic, and finally chaotic rhythmic patterns [68]. As excitability increased, the disruption to eupneic breathing would result in impaired gas exchange *in vivo*. Thus, there is precendent for increasing excitability in the respiratory network resulting in a kind of “depolarization blockade” of normal breathing and a cessation of gas exchange that then results in a precipitous fall in *P*_a_O_2_. We described experiments related to this concept in [17]. Because we have previously shown these transitions are gradual and occur over a wide range of excitability changes, it makes sense to assume that there may be a more gradual progression of the “respiratory collapse”, but we do not yet have clinical data showing how the collapse evolves to the point of need for ventilatory support.

In conclusion, we call for data to be collected on hematocrit in COVID-19 patients, and testing of metabolism and mitochondrial function. Our model predicts changes in oxygen handling and metabolism in silent hypoxemia patients. In addition, we believe the following measures may have untapped predictive value: minute ventilation, oxygen saturation, and breathing frequency. We speculate that some combination of these quantities, if measured on entry to the ER, could help predict the need for ventilator support in the subsequent 48 hours. We also note that there is a need for incorporating oxygen handling dynamics into more sophisticated state-of-the-art respiratory control models, most of which currently focus on CO_2_ and hypercapnea [30]. Finally, a full dynamical systems analysis of why increasing the concentration of hemoglobin shifts the collapse point to lower metabolic demand values in the 7D-O2 model is warranted.

## Methods

Here we provide the equations for the 7D-O2 model introduced in [17].

### Central Pattern Generator (CPG)

A variety of models have been proposed for the central neural circuits generating breathing rhythms, ranging from group-pacemaker networks to individual pacemaker models, and beyond. Here, we adopt the original Butera-Rinzel-Smith (BRS) model (referred to as “model 1” in [18]) proposed as a mechanism for bursting pacemaker neurons in the preBötzinger complex. For simplicity, we represent the CPG with a single BRS unit. Thus our CPG is described by a membrane potential *V* together with dynamical gating variables *n* (a delayed rectifier potassium (*I*_K_) channel activation) and *h* (persistent sodium (*I*_NaP_) channel inactivation). We set two “instantaneous” gating variables *p*_*∞*_ (*I*_NaP_ activation) and *m*_*∞*_ (fast sodium (*I*_Na_) activation) to be equal to their voltage-dependent asymptotic values. We set the *I*_Na_ inactivation gate to be equal to (1 *− n*). The model also includes a leak current (*I*_L_) and a tonic excitatory (*I*_tonic_) current. In summary:

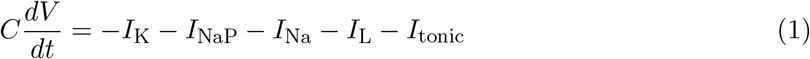

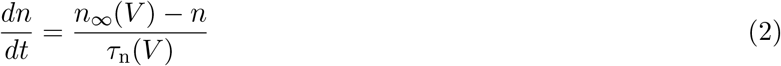

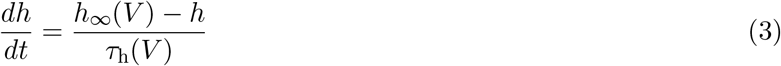

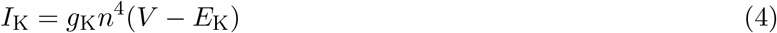

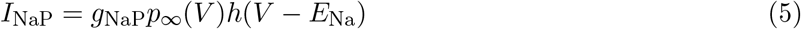

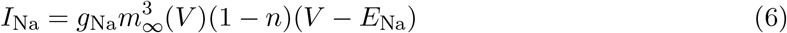

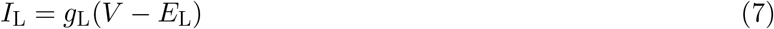

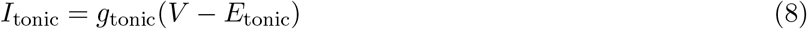

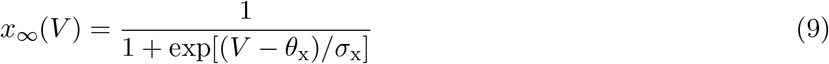

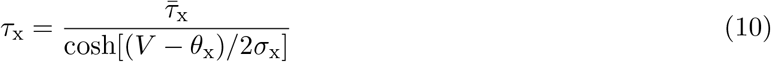

with parameters *C* = 21 pF, *g*_K_ = 11.2 nS, *g*_NaP_ = 2.8 nS, *g*_Na_ = 28 nS, *g*_L_ = 2.8 nS, *E*_K_ = −85 mV, *E*_Na_ = 50 mV, *E*_L_ = −65 mV, *E*_tonic_ = 0 mV, *θ*_n_ = −29 mV, *σ*_n_ = −4 mV, *θ*_p_ = −40 mV, *σ*_p_ = −6 mV, *θ*_h_ = −48 mV, *σ*_h_ = 6 mV, *θ*_m_ = −34 mV, *σ*_m_ = −5 mV, 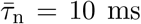, and 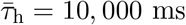.

### Motor pool activity

The output of the CPG is the BRS cell’s membrane potential (*V*), which drives the respiratory muscles through synaptic activation of a motor unit (*α*):

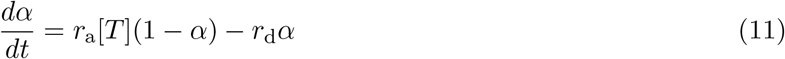

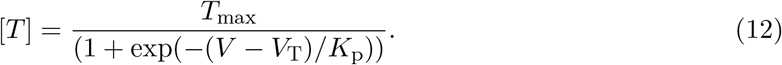

Here, *r*_a_ = *r*_d_ = 0.001 mM^−1^ ms^−1^ sets the rise and decay rate of the synaptic conductance. Also, [*T*] represents the neurotransmitter concentration, with parameters *T*_max_ = 1 mM, *V*_T_ = 2 mV, and *K*_p_ = 5 mV [69].

### Lung volume

The output of the motor unit determines the rise and fall of lung volume (vol_L_):

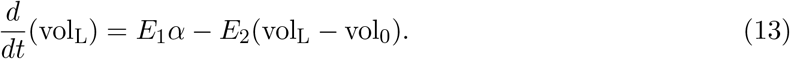

Here vol_0_ = 2 L is the volume of the unloaded lung, and parameters *E*_1_ = 0.4 L and *E*_2_ = 0.0025 ms^−1^ were chosen so that the lung expansion would remain in a physiologically reasonable range [37]. We note that while the low-frequency input of the envelope of CPG burst activity effectively drives changes in lung volume, high-frequency input (such as tonic spiking) does not drive the lung biomechanics effectively. This low-pass filter behavior of the respiratory musculature is analogous to tetanic muscle contraction that occurs in response to high frequency stimulation of motor nerves [70].

### Lung oxygen

At standard atmospheric pressure (760 mmHg), external air with 21% oxygen content will register a partial pressure of oxygen of *P*_ext_O_2_ = 149.7 mmHg. As the lungs expand 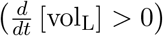, they draw in external air. Our model makes the simplifying assumption that this fresh air mixes instantaneously with the air already present in the lungs. Therefore, the partial pressure of oxygen in the lung alveoli (*P*_A_O_2_) increases at a rate given by the pressure difference between external and internal air, and by the lung volume. In contrast, during exhalation 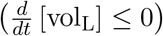, no external air enters the lungs, so the mixing of air stops. During both contraction and expansion of the lung, oxygen moves between the lungs and the blood. The flux of oxygen from the lungs to the blood occurs at a rate determined by the time constant *τ*_*LB*_ = 500 ms, and by the difference in partial pressure of O_2_ between the lungs (*P*_A_O_2_) and the arterial blood (*P*_a_O_2_). The rate of change in *P*_a_O_2_ is thus given by:

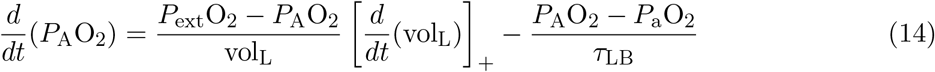

where the notation [*x*]_+_ indicates max(*x*, 0).

### Blood oxygen

To represent the change in *P*_a_O_2_, we write

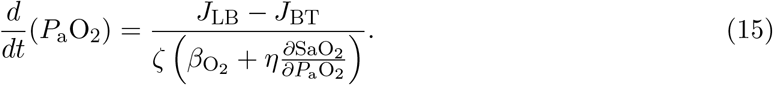

Note the fluxes of oxygen from the blood to the tissues (*J*_LB_) and from the lungs to the blood (*J*_LB_) have units of moles of O_2_ per millisecond. The denominator converts changes in the number of moles of O_2_ in the blood to changes in *P*_a_O_2_. To calculate the flux *J*_LB_, we use the ideal gas law *PV* = *nRT*, where *n* is the number of moles of O_2_, *R* = 62.364 L mmHg K^−1^ mol^−1^ is the universal gas constant, and *T* = 310 K is temperature. The resulting flux depends on the difference in oxygen partial pressure between the lungs and the blood:

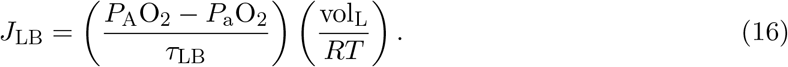

Note that the term *J*_BT_ accounts for both dissolved and hemoglobin-bound oxygen in the blood:

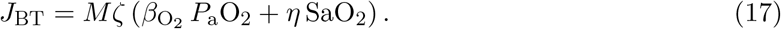

Following Henry’s law, we take the concentration of dissolved oxygen in the blood to be directly proportional to *P*_a_O_2_. The blood solubility coefficient, 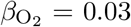 ml O_2_ × L blood^−1^ mmHg^−1^ for blood at 37 degrees C, is the constant of proportionality. The amount of dissolved O_2_ at physiological partial pressures (*P*_a_O_2_ ≈ 80 − 110 mmHg) is insufficient to satisfy the body’s metabolic demand for oxygen. Therefore, most of the blood’s stored oxygen is bound to hemoglobin (Hb).

Cooperative binding of oxygen to the four binding sites in each hemoglobin molecule leads to a sigmoidal hemoglobin saturation curve SaO_2_:

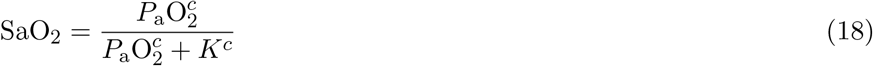

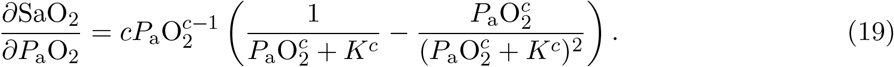

Here, we take the phenomenological parameters *K* = 26 mmHg and *c* = 2.5 from [71].

Our model includes a parameter *M* in Eqn. (17) to capture the rate of metabolic demand for oxygen from the tissues, in units of ms^−1^. Equations (15) and (17) include conversion factors *ζ* and *η* that depend on the concentration of hemoglobin, [Hb] = 150 gm L^−1^, as well as the volume of blood, vol_B_ = 5 L, respectively. The model assumes a molar oxygen volume of 22.4 L. We assume that each fully saturated hemoglobin molecule carries 1.36 ml of O_2_ per gram:

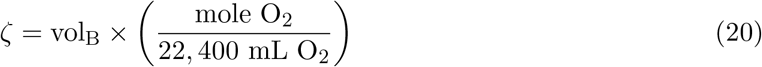

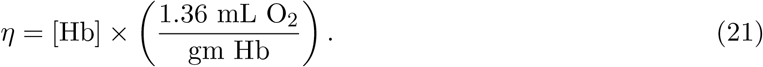

### Chemosensation

Chemosensory feedback from pheripheral chemoreceptors in the carotid bodies, carried to brainstem respiratory circuits via the carotid sinus nerve, close the control loop in our model. These receptors detect reductions in *P*_a_O_2_ and drive the central rhythm generator, as described in more detail in [17]. We model the nonlinear relationship between carotid chemosensory nerve fiber activity and *P*_a_O_2_ as a sigmoidal saturing function, with the firing rate low until *P*_a_O_2_ is reduced below a threshold (normally about 100 mm Hg) and then steep firing rate increases as *P*_a_O_2_ is reduced further [72, 37]. We capture this behavior in our model as a sigmoidal function connecting *P*_a_O_2_ with the conductance representing external drive to the CPG (*g*_tonic_):

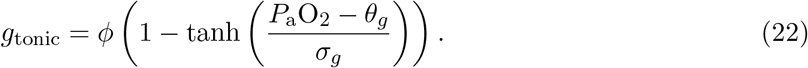

Here, *φ* = 0.3 nS, *θ*_*g*_ = 85 mmHg, and *σ*_*g*_ = 30 mmHg. This conductance closes the control loop in our respiratory control model, since *I*_tonic_ = *g*_tonic_(*V − E*_tonic_) is a term in the CPG voltage equation (1).

We numerically integrated the preceding equations using a variable-order, variable-step stiff solver (`ode15s in MATLAB`).

## Acknowledgments

This work was supported in part by the National Institutes of Health (grants RF1 NS118606-01 and RO1 AT011691-01), the National Science Foundation (grants DMS-2052109, DMS-1555237, and DMS-2152115), and the Oberlin College Department of Mathematics.

## Appendix

**Figure 5:**
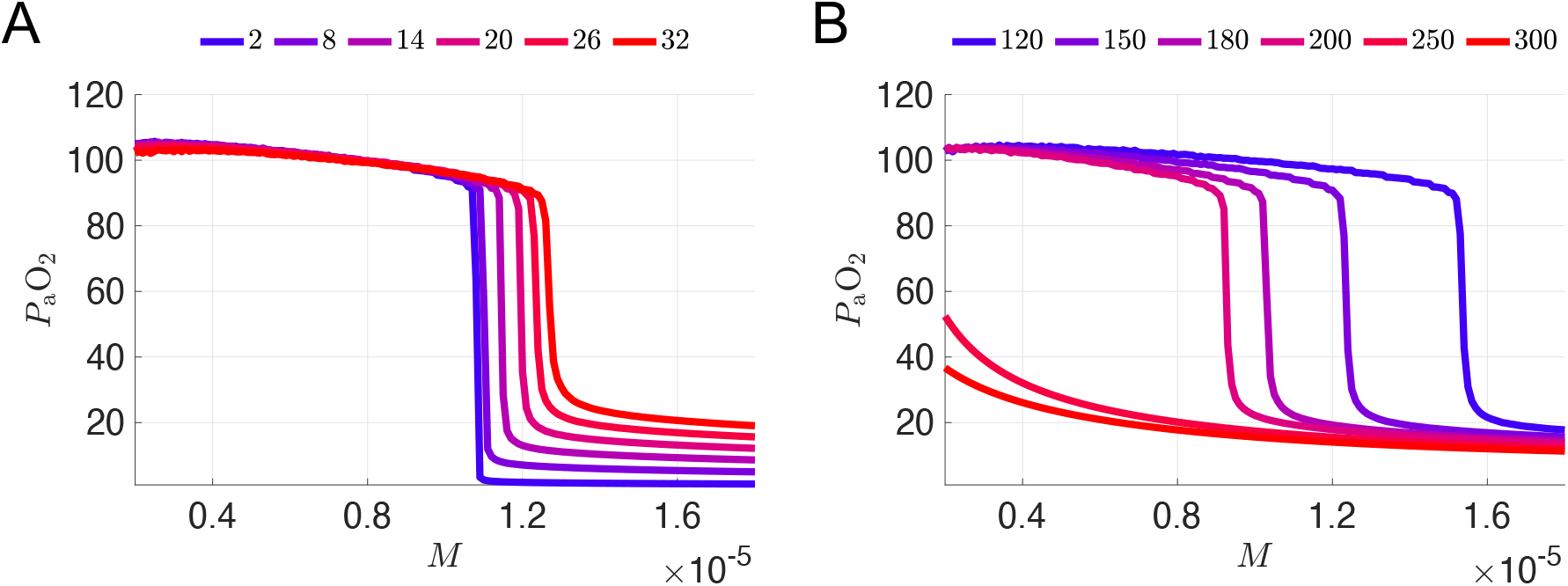
Sensitivity of *P*_a_O_2_ vs *M* curves to variation of hemoglobin parameters. In both panels, all parameters are set to their values from the original 7D-O2 model except for the parameter being varied. **A:** *P*_a_O_2_ vs *M* curves for various hemoglobin binding affinities *K* (including the original value *K* = 26 mmHg). Color scale maps the lowest and highest maximum *P*_a_O_2_ values to blue and red, respectively. **B:** *P*_a_O_2_ vs *M* curves for 6 different values of hemoglobin concentration [Hb] (including the original value [Hb] = 250 gm L^−1^) with the same color scale as in (**A**).

**Figure 6:**
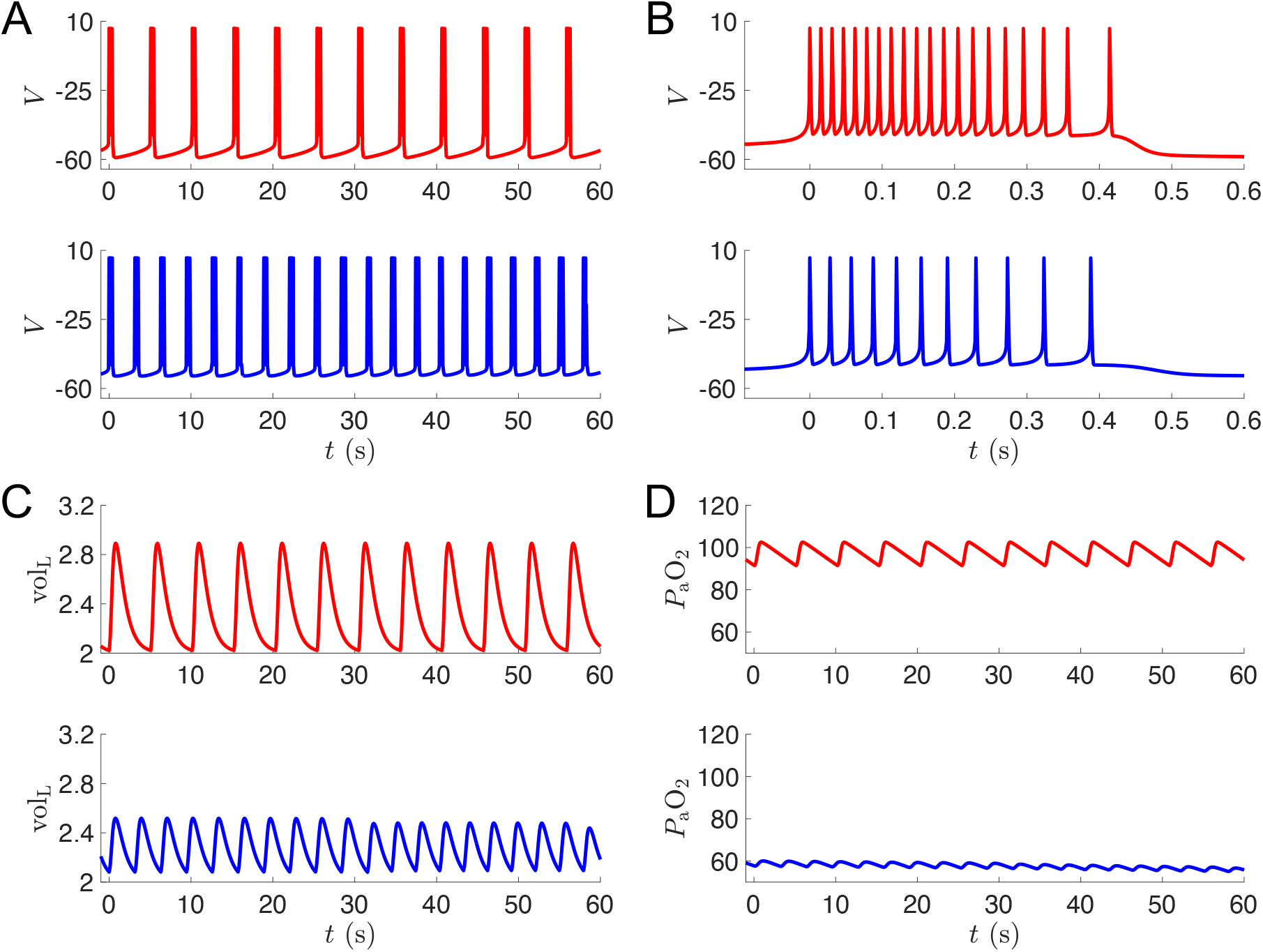
Simulations of putative silent hypoxemia model at high metabolic demand. **A-F:** Output from simulations of the normoxia model (the original 7D-O2 model) and the silent hypoxemia model ([Hb] = 250 gm L^−1^ curve from Fig. 3D) shown in red and blue, respectively, for *M* = 0.97 × 10^−5^ ms^−1^. Voltage traces showing multiple bursts (**A**) and zooming in on a single burst (**B**); lung volume (**C**) and blood oxygen (**D**) traces across multiple bursts.

## Footnotes

1 Model code is available in *ModelDB* at: https://senselab.med.yale.edu/ModelDB/ShowModel?model=229640

